# Application of machine learning in a rodent malaria model for rapid, accurate, and consistent parasite counts

**DOI:** 10.1101/2024.06.05.597554

**Authors:** Sean Yanik, Hang Yu, Nattawat Chaiyawong, Opeoluwa Adewale-Fasoro, Luciana Ribeiro Dinis, Ravi Kumar Narayanasamy, Elizabeth C. Lee, Ariel Lubonja, Bowen Li, Stefan Jaeger, Prakash Srinivasan

## Abstract

Rodent malaria models serve as important preclinical antimalarial and vaccine testing tools. Evaluating treatment outcomes in these models often requires manually counting parasite-infected red blood cells (iRBCs), a time-consuming process, which can be inconsistent between individuals and labs. We have developed an easy-to-use machine learning (ML)-based software, Malaria Screener R, to expedite and standardize such studies by automating the counting of *Plasmodium* iRBCs in rodents. This software can process Giemsa-stained blood smear images captured by any camera-equipped microscope. It features an intuitive graphical user interface that facilitates image processing and visualization of the results. The software has been developed as a desktop application that processes images on standard Windows and Mac OS computers. A previous ML model created by the authors designed to count *P. falciparum*-infected human RBCs did not perform well counting *Plasmodium*-infected mouse RBCs. We leveraged that model by loading the pre-trained weights and training the algorithm with newly collected data to target *P. yoelii* and *P. berghei* mouse iRBCs. This new model reliably measured both *P. yoelii* and *P. berghei* parasitemia (R^2^ = 0.9916). Additional rounds of training data to incorporate variances due to length of Giemsa staining, microscopes etc, have produced a generalizable model, meeting WHO Competency Level 1 for the sub-category of parasite counting using independent microscopes. Reliable, automated analyses of blood-stage parasitemia will facilitate rapid and consistent evaluation of novel vaccines and antimalarials across labs in an easily accessible *in vivo* malaria model.

## Introduction

Eradication of malaria remains a global health priority, with 247 million cases and 619,000 deaths in 2021 (1). Despite the scope of this problem, there is not yet a highly effective vaccine for malaria (2–4). The last few years have seen a drastic spike in malaria cases (1), and resistance is quickly spreading against the most effective therapeutics (5–7). Rapid population growth in malaria-endemic countries, climate-driven epidemiological shifts in Africa (8), and the sudden reappearance of malaria in the American South (9) provide further urgency for finding novel malaria treatments.

Research in blood-stage malaria has an essential and growing role in developing malaria vaccines and therapeutics disease eradication. There is an urgent need for a blood-stage vaccine to reduce clinical malaria symptoms. Furthermore, the spread of artemisinin resistance has accelerated the need for novel antimalarials (5–7). Additionally, monoclonal antibodies capable of neutralizing blood-stage malaria are receiving heightened interest (10–13). *In vivo* rodent malaria models are likely to have a critical role in the development and evaluation of such new therapeutic tools.

The gold standard for analyzing rodent models of malaria involves manual counting of parasites in Giemsa-stained thin blood smears. However, this process is both time intensive and prone to human error. Alternative methods including flow cytometry detection of nuclear stained iRBCs or use of parasites strains expressing luciferase have been reported (14). However, such approaches require specialized equipment, significant optimization and maybe limited to specific parasites strains (eg., parasite expressing fluorescent protein or luciferase). Additionally, they are incapable of collecting the wealth of information found on a blood smear such as infection within reticulocytes vs normocytes and the effect on the stage of the parasite (eg., gametocytes). Additionally, counting parasitemia is well suited for automation as it requires only common laboratory equipment and can significantly reduce the time spent determining parasitemia.

Several groups have previously sought to automate the analysis of human blood smears for malaria (15–22). Yu et al. developed an on-device smartphone-based system (15, 23). It featured a mobile app, Malaria Screener, that was able process the images captured through the eyepiece of a microscope and gave patient-level diagnosis. Another group developed a fully automated end-to-end system, EasyScan Go (24, 25). It’s a digital malaria microscopy device that’s able to take a blood film slide, analyzes it and outputs its diagnosis all at once. Liu et al developed an AI-based system, AIDMAN (16, 17). The findings of their research showed that deep learning models combined with image processing methods can detect and classify parasites with high accuracy.

However, human malaria species, such as *P. falciparum,* do not infect rodent RBC and such ML models may not be applicable to detection of rodent malaria. Three distinct *Plasmodium* species, *P. yoelii*, *P. berghei*, and *P. chabaudi,* which are evolutionarily related to *P. falciparum*, infect rodents. ML researchers have historically focused on human malaria instead of rodents. Only two publications were found that focused on rodent malaria models. Ma et al. proposed a non-deep learning method (26). It did not recognize the morphology of parasites, instead, it only used the color of the Giemsa staining (26). Poostchi et al. also proposed a non-deep learning method, trained on *P. falciparum* iRBCs, and applied it to both human and mouse malaria with only modest success against the latter (27). The large functional conservation of proteins between *P. falciparum* and the rodent malaria species and availability of efficient tools for genetic manipulation makes them an attractive model for *in vivo* evaluation. However, morphological differences in both host and parasites present difficulties when directly utilizing deep learning models trained to identify *P. falciparum*-infected human RBCs for rodent malaria models.

This work used a deep learning model pretrained for *P. falciparum-*infected human RBCs (15) and fine-tuned it on 826 images of *P. berghei* and *P. yoelii*-infected RBCs from mice to developed a ML counting tool for *P. yoelii* and *P. berghei,* achieving an average relative error of 10.74% and 8.31%, respectively. To account for expected domain shifts between laboratories, we retrained the software to account for possible variations due to staining time, microscope, objective, image acquisition platforms etc, to produce a more generalizable model meeting WHO Competency Level 1 for parasite counting (1). Finally, we developed a Windows and MacOS compatible desktop application, Malaria Screener R, which embedded the developed deep learning model. This application lets users process blood smear images in batches, and present detection results with color labels drawn on the images. It also provides a convenient manual correction feature for the mislabeled cells and saves the results to an excel spreadsheet with graphical representations. Overall, we show that this platform can greatly enhance the capacity to efficiently and accurately evaluate preclinical malaria vaccine candidates and therapeutics across different laboratories.

## Materials and Methods

### Overview

In this project, we developed a system to automate the counting of *Plasmodium*-infected RBCs in rodents by using a deep learning-based algorithm. The YOLOv5 object detection model was used for this task. A software application, Malaria Screener R, was developed and used to process blood smear images, review the processed images with overlayed detection results, and export the parasitemia measurements in an Excel ™ file.

The software is designed for use with any camera-equipped microscope. Typically, images captured with these microscopes are stored on a connected computer. For convenience, it is recommended to install Malaria Screener R on the same computer to eliminate the need for image transfer. Once images are captured, users should open the software and select the folder where the images are stored. The software will then process the images accordingly.

On average, our software was able to process one image in 2-3 seconds on one testing machine (2015 MacBook Pro; Processor: 2.7GHz Dual-core Intel Core i5; Memory: 8GB 1867MHz DDR3). Since our software typically needs about 3 images to achieve a good parasitemia estimation, this means it would take under 10s to finish one slide. Such performance can be achieved or surpassed by MacOS and Windows machines with similar or better computing power.

### Software Development

Malaria Screener R was built on top of an existing software called LabelImg. LabelImg is a software commonly used by the deep learning community for data labeling, a process where a human prepares the raw data for training by marking images with the ground truth information. While retaining the valuable features of LabelImg, such as image browsing and image annotation, additional functionalities were incorporated. The new data selection function can handle folders with multi-layer structures which allows a large dataset with multi-layer subfolders to be processed in a one-click fashion. The software will also use folder structure information to organize the results. An image analysis function was added using the aforementioned deep learning model. Furthermore, a user correction function allows quick rectification of model errors, such as mislabeled RBCs, during image browsing. Finally, results display function was added to showcase processed image outcomes, with the option to export results to an Excel file where graphical representations are available. For example, if the dataset contains the data of multiple mice, one plot will have a bar graph to show the estimated percent parasitemia for each mouse. A user manual for software usage is provided in the supplementary data (Supp Figure 1A-B). This software was written in Python. The deep learning model was trained using the Python-based PyTorch framework.

**Figure 1:**
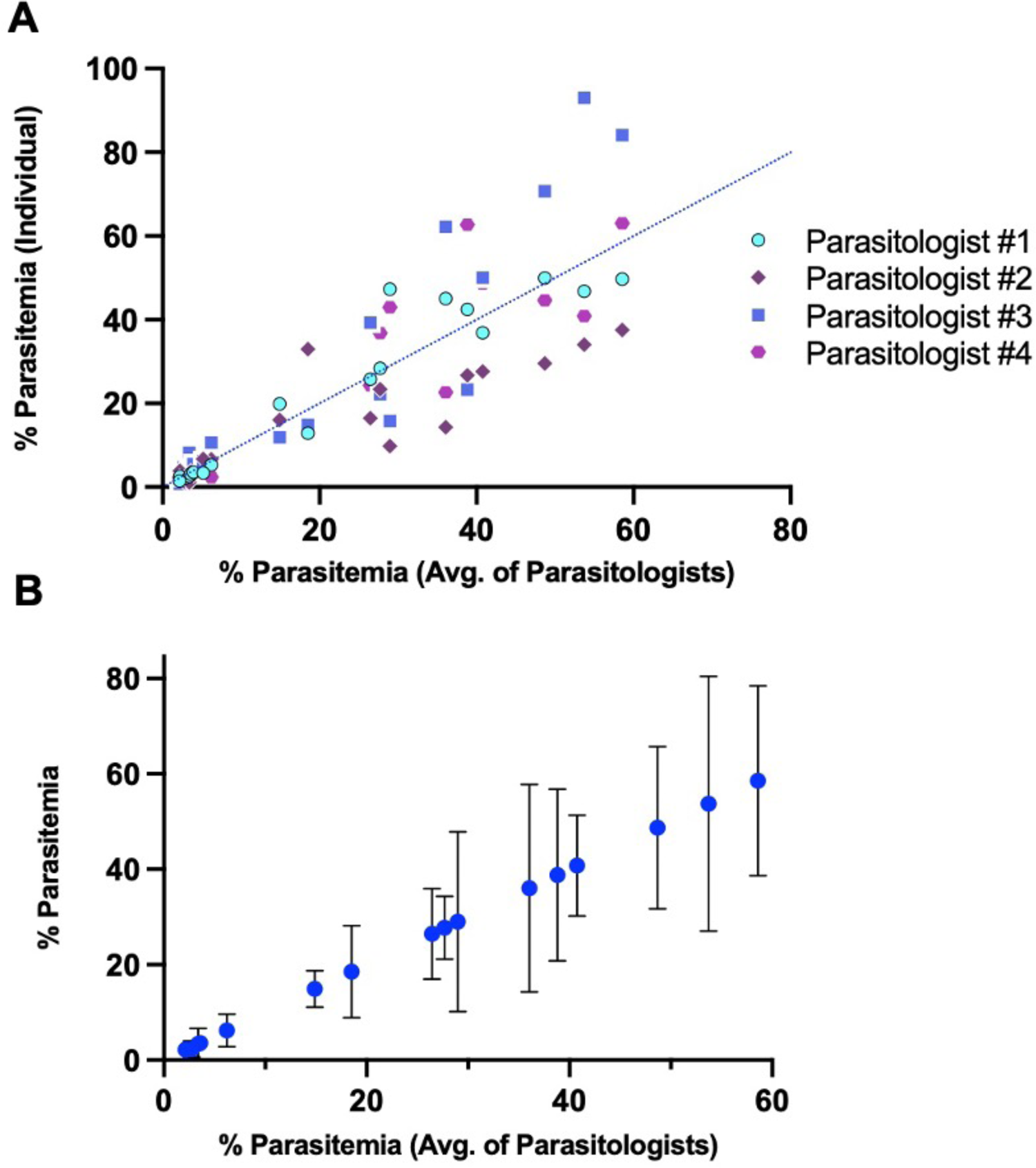
Parasitemia estimates from blood smears are highly variable. A) Estimates of percent parasitemia from individual parasitologists are shown as single points (y-axis) plotted against the average parasitemia measurement of all 4 parasitologists (x-axis). Dotted blue line represents the position of parasitemia measurements if equivalent to manual counts. B) Blue circles represent the percent parasitemia of a single mouse at a single time point (y-axis). Error bars represent standard deviation.

### Model Training with YOLOv5

YOLOv5 is a deep learning-based framework that performs object detection by processing the entire image in a single forward pass through the neural network, making it suitable for real-time applications such as the software application described in this work. YOLOv5 contains five different versions: YOLOv5n, YOLOv5s, YOLOv5m, YOLOv5l and YOLOv5x. The main difference among these models is the amount of feature extraction modules. We chose the YOLOv5l version which gave us a good balance between inference time and detection accuracy.

We used the hyperparameter evolution method provided in the YOLOv5 codebase to help choose the best hyperparameters for training. Table 1 shows the hyperparameters used for training the rodent models in this paper. In addition, original images were resized to 1024×1024 (YOLOv5 requires images to be resized into a square shape), and batch size was set to 6 considering the memory space available on the GPU cards. Model training was performed with a combination of GPU graphics cards including NVIDIA Tesla P100 and V100 models.

**Table 1:** Hyperparameters used for retraining.

| lr0 | lrf | momentum | weight decay | warmup epochs | warmup momentum | warmup bias lr | box | cls | cls_pw |
| --- | --- | --- | --- | --- | --- | --- | --- | --- | --- |
| 0.00285 | 0.396 | 0.98 | 0.00054 | 3.09 | 0.826 | 0.0308 | 0.0515 | 0.966 | 0.908 |
| obj | obj_pw | lou_t | anchor_t | anchors | fl_gamma | hsv_h | hsv_s | hsv_v | degrees |
| 0.2 | 1.22 | 0.2 | 3.9 | 2.0 | 0.0 | 0.0108 | 0.288 | 0.167 | 0.114 |
| translate | scale | shear | perspective | flipud | fliplr | mosaic | mixup | copy_paste |  |
| 0.0461 | 0.219 | 0.0218 | 0.0 | 0.315 | 0.6 | 0.427 | 0.0 | 0.0 |  |

### Training and Testing Model_Rodent_RBCs using images of *P. yoelii*-infected RBCs

Deep learning is a data-intensive approach, requiring a large amount of human annotated data to achieve good results. Therefore, pretrained models are often used to save time and resources. A pretrained model is one that has been trained on large datasets. It can be used as a base model to be fine-tuned for a different but similar task. This will give the benefits of not needing as much data as training a model from scratch. In this study, a model for detecting *P. falciparum* malaria in humans, which we termed Model_Human_RBCs, was used as the pretrained model. This model was developed using a dataset with 965 images from 200 patients collected in Bangladesh. Human and rodent parasite species have similar features. Therefore, the pretrained model for human malaria may be converted for rodent malaria following fine-tuning with additional training images from rodents. An additional 826 images were collected from 10 mice, all infected with *P. yoelii*, and used as training data for the new model, Model_Rodent_RBCs.

Training data was collected in the following manner. Blood smears were made from mouse tail bleeds, which were fixed and then stained with Giemsa (Sigma-Aldrich) (32884-1L) for 10 minutes. Images were taken of blood smears on a Leica ICC50W microscope with a built-in camera using a 100x objective. Images were saved as tiff files in RGB format with dimensions of 2592 x 1944 pixels. One parasitologist annotated all images using the Python application, LabelImg (28). All cells were labeled as infected and uninfected, while nonspecific stains were labeled as debris.

For testing, images of *P. yoelii* and *P. berghei* were compared to manual counts of images. To test Model_Rodent_RBCs’ labeling of *P. yoelii*, 250 images from 9 different mice and 26 unique blood smears were used. To test Model_Rodent_RBCs labeling of *P. berghei*, 43 images from 4 different mice and 11 unique blood smears were used. Enough images were taken for each smear to count at least 500 total RBCs for each unique smear (3-10 images per smear). No mice used in the test set were used in the training set. No user corrections were done following the automated analysis of the data.

### Training and Testing Model_Rodent_RBCs_>10min using darkly stained images

Increased staining time causes darker nonspecific staining and a higher chance of false positives. To improve the accuracy for such images, 100 additional images were collected to retrain the model. All images came from blood smears of *P. berghei* infected mice. Training data was collected and annotated in the manner described above with the following modification: blood smears were stained with Giemsa for either 10, 20, or 70 minutes. The longer stained images were mixed into the training data together with the images from the 10 mice of *P. yoelii.* Training was repeated using the same pretrained *P. falciparum* model, Model_Human_RBCs. Training was done in the same manner as above Model_Rodent_RBCs. However, the original 826 images plus the new 100 images were used to train this model, which we termed Model_Rodent_RBCs_>10min.

For testing, blood smears were taken from *P. berghei* mice and allowed to stain in Giemsa stain (Sigma-Aldrich) (32884-1L) for 10m, 30m or 50m before viewing. A total of 20 smears were analyzed (n = 7, 7, & 6 for 10m, 30m, and 50m groups respectively). The same smears were used for analyses by both Model_Rodent_RBCs and Model_Rodent_RBCs_>10min. Enough images were taken to gather > 500 total RBCs for unique blood smear. No mice used in the test set were used in the training set. No user corrections were done following the automated analysis of the data. Manual counting of images was used as a reference for comparing the accuracy of the two automated models, Model_Rodent_RBCs and Model_Rodent_RBCs_>10min.

### Training and Testing Model_Rodent_RBCs_New_Micro using images from various microscopy platforms

To make the model more generalizable in real-world settings, 99 additional images were collected from three different microscopy platforms to retrain the model, creating Model_Rodent_RBCs_New_Micro. Fifty-four images came from blood smears of *P. berghei*-infected mice, and forty-five images came from blood smears of P. yoelii-infected mice. Training data was collected on a Nikon E800 microscope with a Spot RT Slider top mounted camera using a 100x objective and on a Nikon E600 microscope with a DS-Ri1 top mounted camera with both a 100x and 40x objective. Blood smears were stained with Giemsa for 10 minutes. All files were saved as tiff files in RGB format. For the Nikon E600 with 40x objective, images were saved in the dimensions of 1280 x 1024 pixels. For the Nikon E600 with 100x objective, images were saved with 1920 x 1440 pixels. Finally for the Nikon E800, images were saved as 1600 x 1200 pixels. Again, these new images were mixed into the training data. Then, the model was retrained by repeating the training setup from the previous model, Model_Rodent_RBCs_>10min. This final updated model created from this training set we have termed Model_Rodent_RBCs_New_Micro.

Testing was performed in the following manner. For the Nikon E600 with 40x objective, automated and manual counts were compared using images from 6 different mice and 10 unique blood smears. A total of 16 images were used for these 10 blood smears. Finally for the Nikon E600 with 100x objective, automated and manual counts were compared using images from 6 different mice and 8 unique blood smears. A total of 77 images were used for these 8 blood smears. Enough images were taken to gather greater than 500 total RBCs per unique blood smear. No mice used in the test set were used in the training set. No user corrections were done following the automated analysis of the data.

### Parasitemia estimates by expert parasitologists

Expert parasitologists consisted of PhD students and post-doctoral fellows, who have completed greater than 3+ years of malaria research. Parasitemia estimates by the 4 expert parasitologists were performed in the following manner. First, the total number of RBCs in one field was counted. Then, the number of infected RBCs in the same field was counted. Next, the stage was moved vertically to the next field, which was assumed to have a similar distribution of RBCs, and the number of infected and total RBCs was counted. This process was then repeated for 10 fields. The number of total RBCs was estimated by averaging the counts from the first and last fields examined, and then multiplying this average by the ten fields counted.

### Manual Counting of individual RBCs in images

The same images uploaded into the automated program were viewed by an expert parasitologist. With the assistance of the application ScreenGrid^TM^, each infected and uninfected RBC within the image was counted. For cells on the border of images, only those cells, where the majority of the cells were present in the image frame, were included in counts. Likewise, cells largely obscured by debris were not included in counts. Infected and uninfected RBCs were counted 2x and the average of the two values was used. If the two counts varied by more than 10%, a third recount was done.

### Infection of mice

Female C57Bl/6 mice (Jackson Laboratory) and female Swiss Webster mice (Charles River Laboratories) of varying ages (4-14 wk) were used in this study. All animal experiments were approved by the Johns Hopkins Animal Care and Use Committee (ACUC), under protocol MO22H289. Mice were infected via the tail vein with 10^4^ infected RBCs, containing either the *P. yoelii* or *P. berghei* parasite species. Mice were euthanized if parasitemia reached 75% or higher.

### Parasite strains

Both non-lethal *P. yoelii* XNL and lethal *P. yoelii* 17XL strains were used for P. *yoelii* infections. *P. berghei* strain ANKA parasites were used for *P. berghei* infection.

### Production and staining of blood smears

Mouse tail bleeds were performed daily starting 2-3 days post infection. A small drop of blood was placed on a glass slide and a smear was made. Smears were placed for a few seconds in 100% methanol for fixation and then in Giemsa stain (Sigma-Aldrich) (32884-1L) for 10 minutes, unless otherwise noted within the study. Slides were allowed to air dry or in some cases dry time was sped up using a hair dryer. Slides were then viewed underneath a light microscope.

## Results

### Blood Smear Estimates of *P. yoelli* and *P. berghei* by parasitologists are highly variable

Four parasitologists were tasked with performing estimates of parasitemia on thin blood smears on the same 19 Giemsa-stained smears of *P. yoelli* and *P. berghei* infected RBCs, 10 *P. berghei* and 9 *P. yoelii.* Each parasitologist used a standard system for counting parasitemia, counting all infected RBCs in each field and estimating the total number of RBCs in each field (detailed further in the methods). Parasitemia varied significantly between parasitologists (Figure 1A & 1B) with an average relative standard deviation of 43.31% among the 19 blood smears tested (Supp Figure 2). Significant deviation was found at all measurable parasitemia levels, and standard deviation showed no correlation with parasitemia levels (R^2^ = 0.0099)(Supp Figure 2).

**Figure 2:**
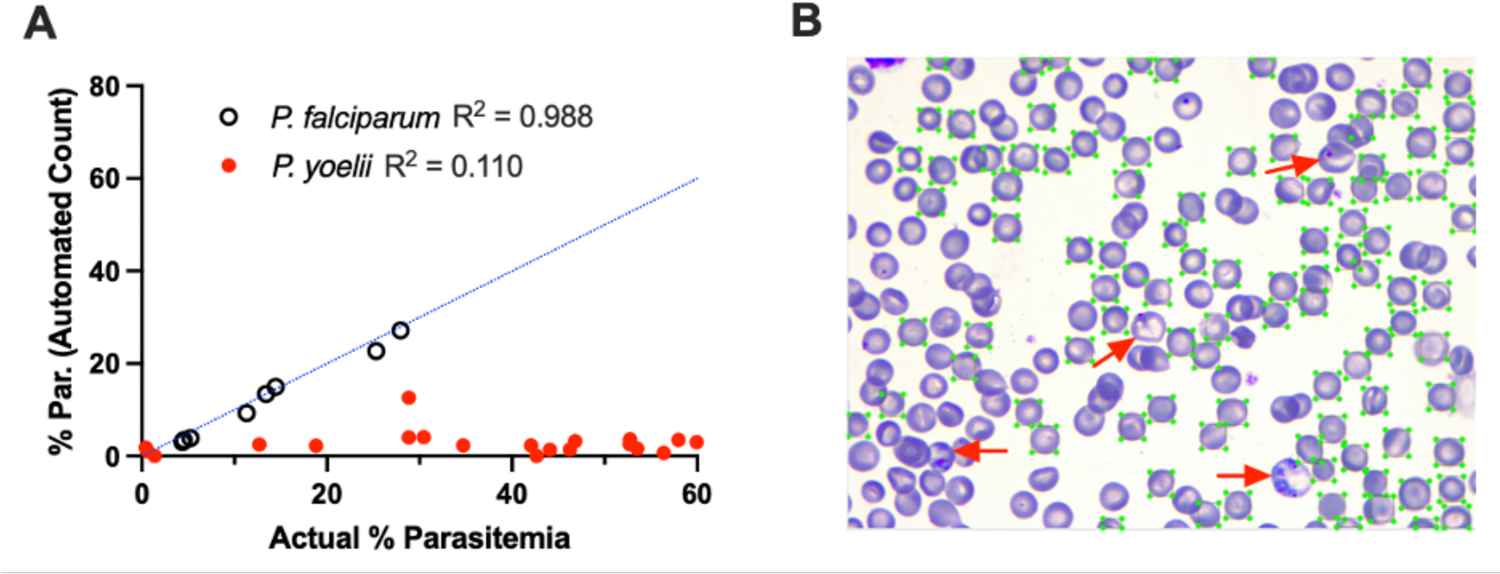
Previous model trained on *P. falciparum*-infected human RBCs fails to detect *P. yoelii*-infected mouse RBCs. A) Each circle represents a parasitemia measurement determined using the previous *P. falciparum*-trained malaria screener model trained on *P. falciparum*-infected RBCs (black open circles) or *P. yoelii*-infected RBCs (red closed circles). X-axis represents percent parasitemia measured by manual counting of RBCs in images by expert parasitologist. Dotted blue line represents the position of parasitemia measurements if equivalent to manual counts. B) Example image of automated detection of RBCs using this model. Green boxes circumscribe all RBCs (both infected and uninfected) recognized by the model. Red arrows indicate infected RBCs not recognized by the software.

### Previous *P. falciparum* trained model fails to accurately measure parasitemia in *P. yoelii-*infected rodent RBCs

Previously, our group developed and optimized a ML model (Model_Human_RBCs) to identify *P. falciparum*-infected human RBCs in Giemsa-stained thin smears from patients in a clinical setting (15). In the current study, this model was applied to measure both *P. falciparum*-infected human RBCs and *P. yoelii*-infected mouse RBCs. While the program accurately measured parasitemia in *P. falciparum*-infected human RBCs, it failed to do so in *P. yoelii*-infected mouse RBCs (Figure 2A). When testing Model_Human_RBCs on *P. falciparum*-infected RBCs, automated counts correlated highly with manual counts (R^2^ = 0.988) (14.25% relative error rate). In contrast when testing Model_Human_RBCs on *P. yoelii*-infected RBCs, automated counts correlated poorly with manual counts (R^2^ = 0.110) (100.83% relative error rate). On the *P. yoelli*-infected RBC test set, the model underperformed in two areas, failing to identify mouse RBCs (both infected and uninfected) and to differentiate infected from uninfected mouse RBCs (Figure 2B).

### Model trained on images of *P. yoelii*-infected RBCs accurately estimates rodent parasites at a wide parasitemia range and RBC densities

Because of the limitations of the previous model trained on *P. falciparum*-infected human RBCs in identifying *P. yoelii*-infected mouse RBCs, we sought to develop a new model capable of labeling such cells, which we termed Model_Rodent_RBCs (Figure 3). We fine-tuned the model using 826 images taken from 10 different *P. yoelii*-infected mice at varying parasitemia levels (Figure 3). This led to a substantial improvement in identification of uninfected and infected RBCs with the automated counts closely matching the manual counts determined from the same set of images (R^2^ =0.9933) (Figure 4A, Supp Figure 3). For parasitemia levels greater than 1 percent, relative error was low between automated and manual counts (mean relative error =10.74%) (Figure 4B Supp Figure 3). To evaluate accuracy, we created two different % parasitemia reference standards (Supp. Figure 3A). The first was based on manual estimates of parasitemia in blood smears. In this method, typically used for parasitemia measurements, the total number of cells in each microscopy field is estimated from a subset of fields (% parasitemia = iRBCs in 10 fields/ RBCs in 2 fields picked at random*5). In the second method, we counted every cell in the images directly used for the test data, a more stringent and time-consuming approach (% parasitemia = total iRBCs/total RBCs). While the former standard helps validate the accuracy in a more real-world setting, the latter provides a more precise manner for determining the accuracy of the model in classifying cells in the images analyzed. Notably, we see improvements in the accuracy of the automated system vs. manually counting when comparing against both standards (Supp. Figure 4A-C). Compared against manual estimates of parasitemia in blood smears, the automated system demonstrated better accuracy than 3 of 4 expert parasitologists and accuracy equivalent to the 4^th^ parasitologist. Compared against individual counting of RBCs in images, the automated system demonstrated better accuracy than all 4 parasitologists.

**Figure 3:**
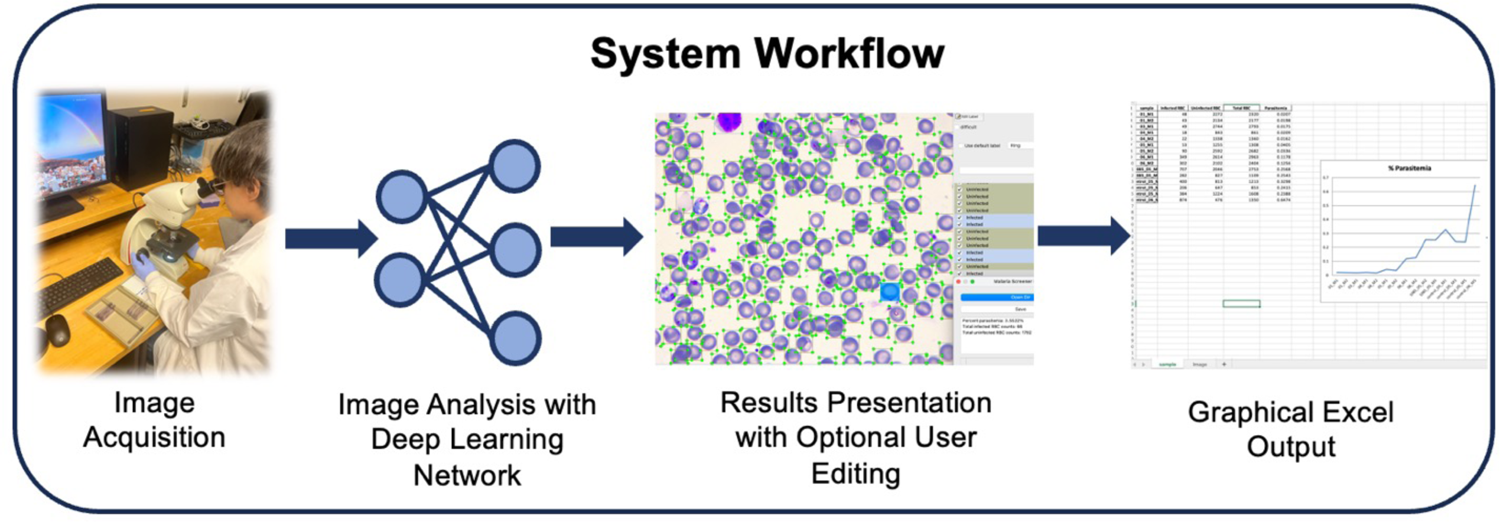
Schematic demonstrating workflow of automated malaria detection system. Images are captured of Giemsa-stained rodent thin blood smears using a light microscope with a built-in camera. Images are uploaded to software, and each cell is detected and then labeled as infected or uninfected using a pre-trained convolutional neural network (CNN). Each annotated image is then displayed in a graphical interface allowing users to verify and -if necessary-correct labels. Upon saving each image, an excel sheet is created with a graphical output of percent parasitemia measurements. Graphical summary of results is organized using the directory structure of uploaded images and user input of groupings upon saving.

**Figure 4:**
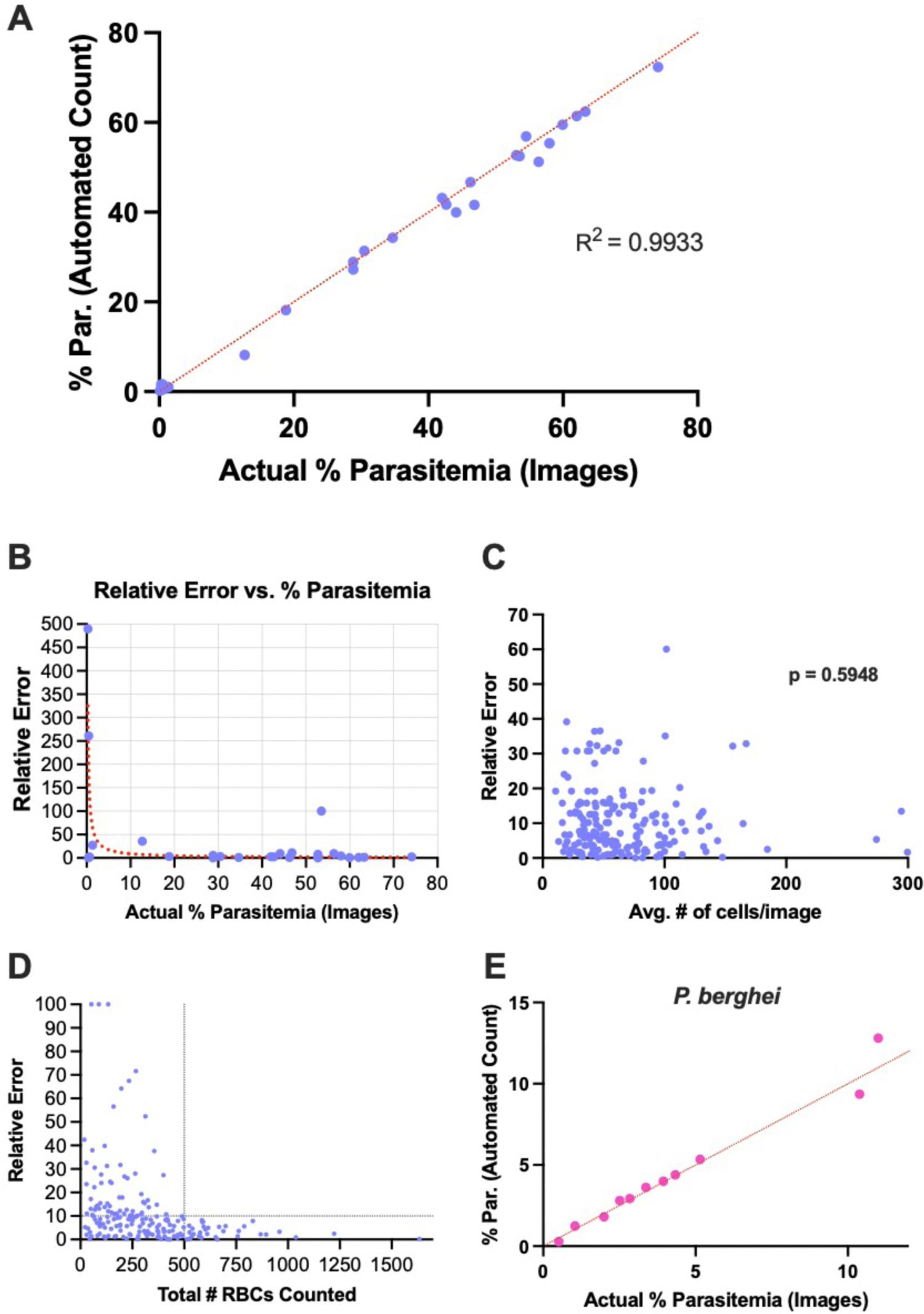
Automated detection method calculates percent parasitemia of *P. yoelii* and *P. berghei-*infected cells with high accuracy. A) Relationship between manual counting of parasitemia (x-axis) and percent parasitemia measured using automated detection method. Each blue dot represents the percent parasitemia of a *P. yoelii* infected mouse at a single time point. Red dotted line indicates position of each point if manual and automated counts are equal. B) Relative error of automated parasitemia measurements of *P. yoelii*-infected mice. Blue dots represent relative error between automated and manual measurements for 1 mouse at 1 time point utilizing Model_Rodent_RBCs. Relative error is calculated as the absolute value of ((% parasitemia automated count – % parasitemia manual count)/(% parasitemia manual count)*100). Red dotted line represents nonlinear fit of relative error vs. % parasitemia. C) Relationship between relative error (y-axis) and RBC density (x-axis). Each blue dot represents the total number of RBCs in a *single* image plotted against the relative error between manual and automated counts for that image. D) Relationship between relative error (y-axis) and total number of RBCs (infected and uninfected) counted by automated software. Blue dots show the total number of cells counted by the software and the relative error compared to manual measurements. Data points are based on parasitemia measurements of one or more images of blood smears from a single mouse at a single time point. Additional images from the same blood smears were added in series. Intersection of black dotted lines indicates stabilization of relative error values around 500 total RBCs counted. E) Pink dots show the relationship between manual and automated parasitemia measurements of *P. berghei-*infected RBCs. Red dotted line shows the position of points if measurements are equal.

At parasitemia levels >1% the program produced highly accurate counts (median relative error = 5.88%). However, at parasitemia levels <1% the error rate was higher (74.78% median relative error) (Figure 4B). At very low parasitemia levels false positives can lead to high error rate. Indeed, the errors observed were typically characterized by the presence of one or more false positive infected RBCs in the automated detections. Our ML software, Malaria Screener R, provides the user the ability to easily correct the label on only a few RBCs in order to correct such false positives and account for these errors.

Overlapping RBCs may often be difficult for automated software to distinguish. In order to test the model’s ability to handle such situations, we analyzed the *P. yoelii*-trained model’s (Model_Rodent_RBCs) performance at a range of cell densities, up to 300 RBCs per field using a Leica DMi8 microscope with an integrated camera.

Model_Rodent_RBCs performed similarly across all tested RBC densities (Figure 4C). There was not a significant relationship between the number of RBCs per image and relative error (Pearson correlation coefficient = 0.03853, p = 0.5948). Importantly, error rates for the program tended to decrease as the number of RBCs analyzed increased. In order to better inform users’ decision of how many images to upload to the software, we analyzed the relationship between relative error of the program and the total number of RBCs analyzed. Relative error between automated and manual counts appeared to stabilize below 10 percent around 450-500 total RBCs counted (Figure 4D). This value corresponds to about 3 images taken containing 150 RBCs per image, a density which we previously demonstrated the program can reliably measure (Figure 4C).

*P. yoelii* and *P. berghei* are two commonly used rodent malaria parasites in pre-clinical studies. As the dataset for fine-tuning the model used only *P. yoelii*-infected RBCs, we tested the program’s ability to measure parasitemia from 11 different blood smears of *P. berghei* infected mice at parasitemia ranging from 0.5 to 11%. The software accurately measured parasitemia across a range of values with a median relative error of 8.33% for parasitemia values >1%, indicating that the program performs comparably on both the rodent malaria parasites (Figure 4E).

### Giemsa stain time of blood smears has a significant effect on model performance

While comparing manual and automated counts, we noticed that the accuracy of the program was affected when different individuals were performing the Giemsa stain of blood smears. Examining this issue further, we found that the difference was related to a change in the amount of time slides were stained with Giemsa. For slides stained 10-15min, the normal recommendation, the program annotated with high accuracy. However, when the stain time was extended to 30-50min, the accuracy of the program decreased drastically, as increased nonspecific staining led to a higher number of falsely labeled infected RBCs (Supp Figure 5A). To account for such potential differences, we trained a separate model with an additional 100 new images from blood smears stained for atypically long times (Model_Rodent_RBCs_>10min). Following retraining, there was a decrease in error rates for blood smears stained for 30min and 50min (Supp Figure 5B). However, error rates for these extended stain times remained much higher than images from smears stained for 10min. Interestingly, accommodating for longer staining protocols also improved the performance of the program at parasitemia below 1% to a median relative error of 17% compared to 35% before optimization (Supp Figure 5C). Therefore, it is recommended not to stain longer than 10 min.

**Figure 5:**
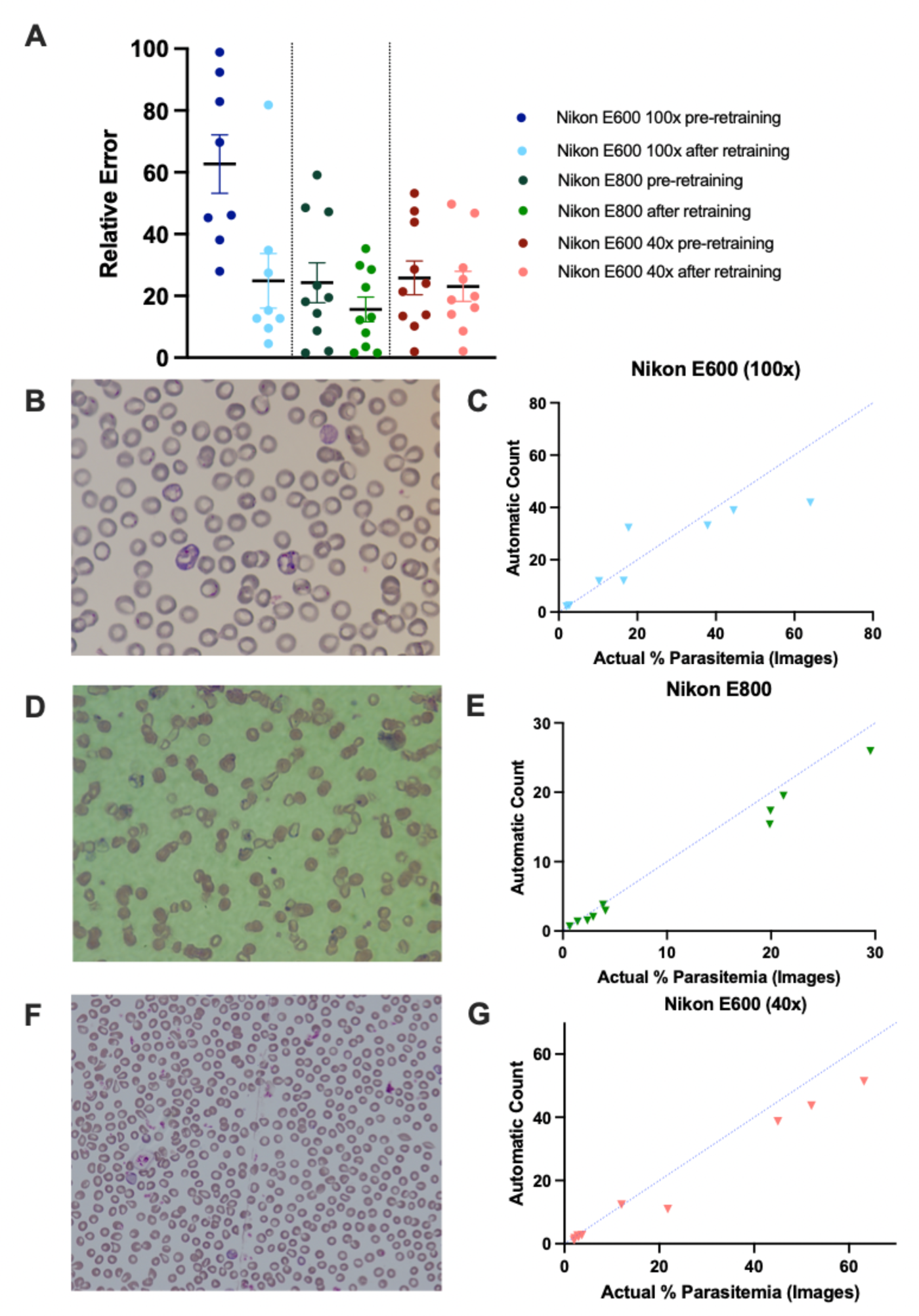
Model is generalizable to images captured on different microscopes A) Relative error (y axis) is shown for the difference between the original model’s annotations and manual counts using images from blood smears on a Nikon E600 with a 100x objective (dark blue, left), on a Nikon E800 with a 100x objective (dark green, center), and on a Nikon E600 with a 40x objective (dark red, right). Relative error was calculated for each mouse at one time point as the absolute value of ((% parasitemia automated count – % parasitemia manual count)/(% parasitemia manual count)*100). The model was then retrained with an additional 99 images total from these microscopes and updated to a single new model, Model_Rodent_RBCs_New_Micro. Relative error was recalculated for each data point and displayed in light blue, light green, and light red for the Nikon E600 (100x objective), Nikon E800 (100x objective), and Nikon E800 (40x objective) respectively. Data is mean ± SEM. Example images of blood smears taken on a Nikon E600 with a 100x objective (B), on a Nikon E800 with a 100x objective (D), and on a Nikon E600 with a 40x objective (F). Relationship between manual counting of parasitemia (x-axis) and percent parasitemia measured with automated detection method, using images taken on a Nikon E600 with a 100x objective (C), on a Nikon E800 with a 100x objective (E), and on a Nikon E600 with a 40x objective (G). Each dot represents the percent parasitemia of a *P. yoelii* or *P. berghei* infected mouse at a single time point. Blue dotted line indicates position of each point if manual and automated counts are equivalent.

### Model is generalizable to different microscopes and image-acquisition platforms

Differences in cameras, microscopes, and objectives used may alter image quality and influence count accuracy. Indeed, images acquired using three different microscopes (Nikon E800 with 100x objective, Nikon E600 with 40x objective, and Nikon E600 with 100x objective) varied significantly in quality and size of RBCs (Figure 5B, 5D, & 5F), and all 3 microscopes showed higher relative error values than the Leica microscope utilized for capturing images used for ML training in Model_Rodent_RBCs. This lower accuracy appears to be due to undercounting in 24 of 27 test cases. To address this domain shift, the model was fine-tuned using 99 additional annotated images collected from different microscopes, 54 images of *P. berghei* parasites and 45 images of *P. yoelii* parasites (Model_Rodent_RBCs_New_Micro). Following fine-tuning, automated counts for all 3 microscopes fit more closely to expected values, determined by manual counting of individual cells in images (Figure 5A). Average relative error rates were 15.64%, 23.07%, and 24.84% for the Nikon E800 with 100x objective, Nikon E600 with 40x objective, and Nikon E600 with 100x objective respectively (Figure 5A, 5C, 5E, 5G). The <25% error rate obtained for all 3 additional microscopes following training optimization reached level 1 competency when measured using WHO guidelines for assessment of microscopists in human malaria diagnosis(29).

### Automated model predicts parasitemia with greater precision than manual counting

Comparison of *in vivo* models of malaria both within and between laboratories requires a consistent means of measuring parasitemia. The large standard deviation of parasitemia measurements between parasitologists counting the same blood smears (Figure 1A & 1B) hinders such comparisons. To evaluate precision of the automated platform vs. manual counting of blood smears, four parasitologists were tasked with measuring the same 11 blood smears using standard manual counting and four users were tasked with measuring the same 11 blood smears using the automated method developed in this study (Figure 3). The automated program with both deep learning networks, Model_Rodent_RBCs and Model_Rodent_RBCs_New_Micro, demonstrated significantly greater precision, measured via relative standard deviation, than parasitologists manually counting blood smears (Figure 6A & 6B). Across all parasitemia levels, the automated program showed greater precision than manual counting (Figure 6A & 6B). Interestingly, the automated method using the updated model, Model_Rodent_RBCs_New_Micro, showed even greater improvement over the original model, Model_Rodent_RBCs.

**Figure 6:**
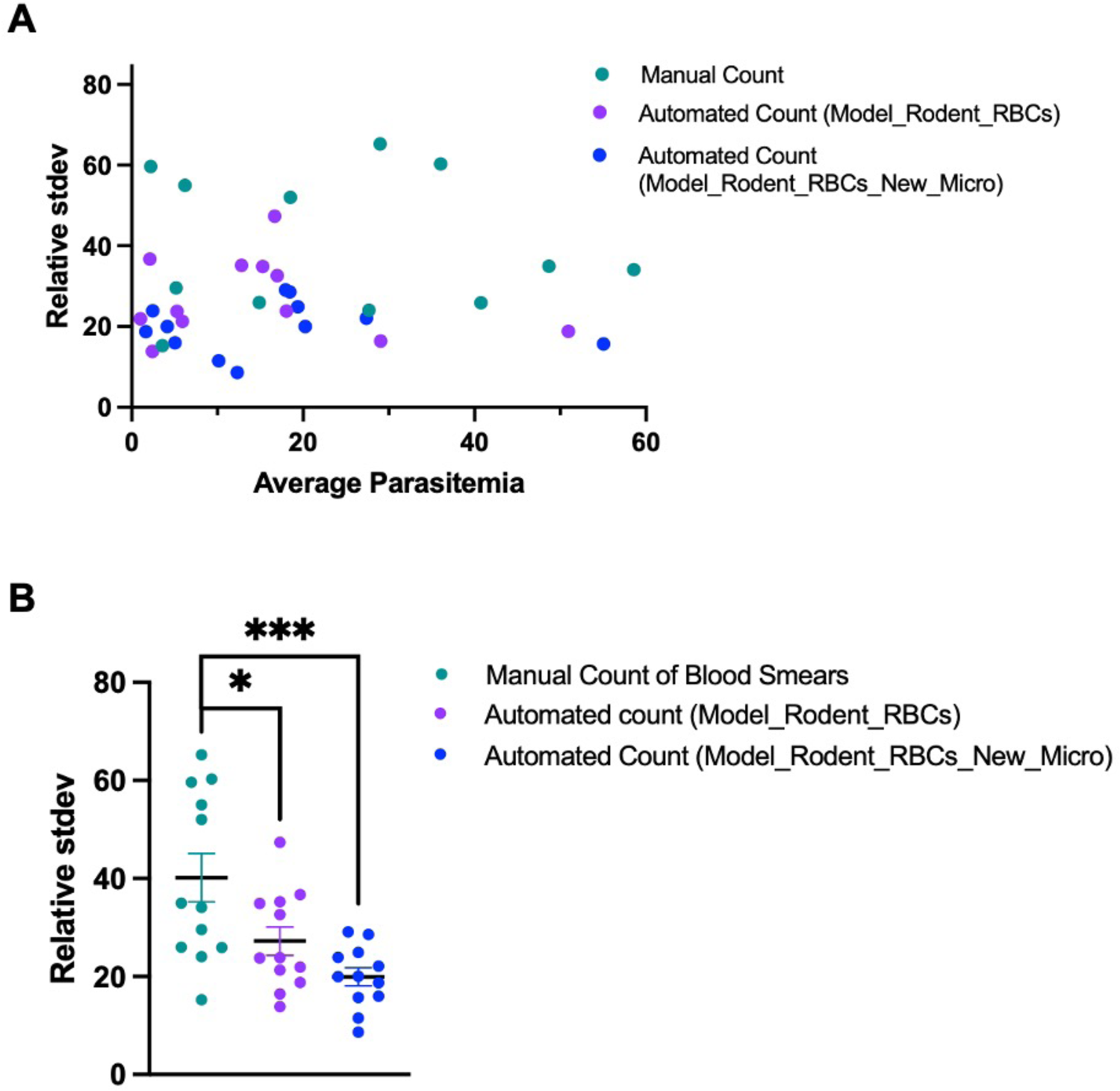
Automated model predicts parasitemia with greater precision than manual counting. A) Relative standard deviation of parasitemia measurements (y-axis) is shown for 4 different expert parasitologists manually counting blood smears (Teal) and 4 parasitologists using the automated software trained on parasites infecting rodent RBCs (Model_Rodent_RBCs = purple, Model_Rodent_RBCs_New_Micro = blue) across a wide range of parasitemia measurements. X-axis values denote the average of measurements by 4 users manually counting blood smears or using the automated software. Relative standard deviation was calculated as (100*Standard deviation/Mean % Parasitemia). B) Relative standard deviation for all parasitemia measurements is shown when 4 users manually count blood smears (teal) vs. when 4 users utilize the automated software (Model_Rodent_RBCs = purple, Model_Rodent_RBCs_New_Micro = blue). P values were calculated using unpaired, two-tailed t tests.

## Discussion

Rodent models of malaria can play a critical role in testing preclinical therapeutics and vaccines for prioritizing subsequent evaluation in non-human primates or humans. However, the gold standard for analyzing blood stage malaria in these models largely rely on counting of blood smears, a time-consuming and repetitive process. This method can also be error prone due to the use of estimates rather than actual counts of total RBCs and parasitologists biases in classifying infected vs. uninfected RBCs. Parasitologists typically estimate the total number of RBCs they are counting as individual counting of RBCs in all fields is too time-consuming. In this study, we find a remarkably high average relative standard deviation in parasitemia measurements of 43.31% between expert parasitologists.

We utilized transfer learning to adapt a machine learning model, which was previously trained on *P. falciparum*-infected human RBCs, to identify *P. yoelii*-infected mouse RBCs. We developed a Windows and MacOS compatible desktop app incorporating this model and capable of providing accurate parasitemia measurements with limited user input (released with publication of the study). This software requires only a microscope with a mounted or built-in camera and a computer for analysis. Using this semi-automated process, the user will make blood smears, collect images on a microscope with the camera, and process the images with the software. The software then rapidly annotates each cell as infected or uninfected and provides both a table and graphical representation of results in a spreadsheet.

In contrast to standard manual counting methods, which count infected RBCs but estimate the total number of RBCs, the automated program counts all uninfected and infected RBCs, theoretically allowing for more accurate measurements. Supporting this theory, the automated system demonstrates better accuracy compared to expert parasitologists’ manual counts of the same smears. This automated program (Model_Rodent_RBCs) demonstrated an average relative error of 10.74% and 8.31% compared to image-based manual counts for *P. yoelii* and *P. berghei*, respectively, across a wide range of RBC densities and parasitemia values (1-75%). Importantly, the model accurately identifies RBCs at very high densities, allowing for acquisition of only a few images to achieve robust sample sizes. In addition to improving accuracy, the model also provides significantly more consistent results. Both counting of all RBCs, as opposed to estimating total RBCs, as well as the elimination of parasitologist biases in parasite classification likely contribute to the improved consistency of the automated program over manual counting. This result, importantly, allows for effective comparison of different therapeutics across distinct rodent malaria studies.

The initial model, Model_Rodent_RBCs, trained using images from a single microscope, demonstrated a loss in accuracy when measuring parasitemia using images taken on different microscope platforms. Interestingly, following retraining with only the addition of a small number of images from different platforms, the model met accuracy levels comparable to the WHO Level 1 standard for counting *P. falciparum* (29). Remarkably, after switching from a 100x to a 40x objective and thereby drastically decreasing the apparent size of both RBCs and parasites, the model did not show a loss of accuracy. These results suggest broad applicability of this model to a variety of microscopes and laboratories.

During this study, we found that the Giemsa stain time of blood smears impacted the accuracy of the program, with longer than normal stain times leading to a significant loss of accuracy. Retraining with additional images from longer stained smears was able to significantly improve the accuracy of measurements on such smears but still demonstrated a loss of accuracy. For this reason, we strongly recommend limiting Giemsa stain times to 10-15 minutes when using this program. Retraining, however, offered an unexpected benefit to the accuracy of the program. Model_Rodent_RBCs_>10min demonstrated an improved ability to distinguish false positives. This improvement is most apparent for images with very low parasitemia values (<1%), where a single false positive may drastically affect the parasitemia and relative error. In Model_Rodent_RBCs_>10min, there was a 4-fold decrease in the average relative error in parasitemia measurements for images below 1% parasitemia, as compared to Model_Rodent_RBCs. While the updated model still only has moderate accuracy below 1% parasitemia (median relative error 16.97%), the software’s built-in function to optionally correct mislabeled RBCs offers a rapid means of correcting measurements at such low parasitemia values. All results in this study do not include optional corrections, and any such corrections will lead to greater improvements in accuracy.

The program developed in this study provides a consistent, accurate and efficient method for the analysis of infected RBCs in rodent malaria models. Importantly, this ML-based automated tool, Malaria Screener R, can provide reliable parasitemia counts and enable fast evaluation of novel vaccines and antimalarials in an easily accessible *in vivo* malaria model. The 1025 image data set utilized in this study has also been fully annotated with the parasite stage (gametocyte, ring, schizont, trophozoite), the number of parasites in each RBC, and the identification of both infected and uninfected immature RBCs (reticulocytes). Future updates to the software may incorporate staging parasites, separate counting of multiply infected RBCs, and identifying infected and uninfected immature RBCs (reticulocytes).

## Acknowledgements

We thank Dr. Jake Ruddy for valuable discussions regarding this study. We also thank Natalie Gobrial for her help annotating uninfected RBCs in this study. We thank Dr. Photini Sinnis and the Sinnis Laboratory for use of their Nikon E600 microscope in this study. This work was supported in part by the Lister Hill National Center for Biomedical Communications of the National Library of Medicine (NLM), National Institutes of Health. This work was also supported by the Johns Hopkins Malaria Research Institute and the Bloomberg Philanthropies.

## Financial Support

S.Y. was supported by the NIH (T32 AI138953). N.C. and O.A-F. were supported by Johns Hopkins Malaria Research Institute. E.C.L. was supported by the NIH (T32AI138953). P.S. was supported in part by the NIH (R01AI155598) and Johns Hopkins Malaria Research Institute.

## Disclosures

None. Software is made freely available. All authors have no current financial interests in the program.

## Software Availability

The software is available for free download at https://lhncbc.nlm.nih.gov/LHC-research/LHC-projects/image-processing/malaria-screener.html

## Authors Further Information

1. Sean Yanik

– Department of Molecular Microbiology and Immunology and Malaria Research Institute, Johns Hopkins School of Public Health, Baltimore, MD, USA 21205
2. Hang Yu

– National Library of Medicine, National Institutes of Health, Bethesda, MD, USA 20894
3. Nattawat Chaiyawong

– Department of Molecular Microbiology and Immunology and Malaria Research Institute, Johns Hopkins School of Public Health, Baltimore, MD, USA 21205
4. Opeoluwa Adewale-Fasoro

– Department of Molecular Microbiology and Immunology and Malaria Research Institute, Johns Hopkins School of Public Health, Baltimore, MD, USA 21205
5. Luciana Ribeiro Dinis

– Department of Molecular Microbiology and Immunology and Malaria Research Institute, Johns Hopkins School of Public Health, Baltimore, MD, USA 21205
6. Ravi Kumar Narayanasamy

– Department of Molecular Microbiology and Immunology and Malaria Research Institute, Johns Hopkins School of Public Health, Baltimore, MD, USA 21205
7. Elizabeth C. Lee

– Department of Molecular Microbiology and Immunology and Malaria Research Institute, Johns Hopkins School of Public Health, Baltimore, MD, USA 21205
8. Ariel Lubonja

– Department of Computer Science, Johns Hopkins Whiting School of Engineering, Baltimore, MD, USA 21218
9. Bowen Li

– Department of Computer Science, Johns Hopkins Whiting School of Engineering, Baltimore, MD, USA 21218
10. Prakash Srinivasan

– Department of Molecular Microbiology and Immunology and Malaria Research Institute, Johns Hopkins School of Public Health, Baltimore, MD, USA 21205
11. Stefan Jaeger

– National Library of Medicine, National Institutes of Health, Bethesda, MD, USA 20894

## Notes

### Competing Interest Statement

The authors have declared no competing interest.

