## Supplementary material for "Application of machine learning in a rodent malaria model for rapid, accurate, and consistent parasite counts": Suppl Figs

**A**

**B**

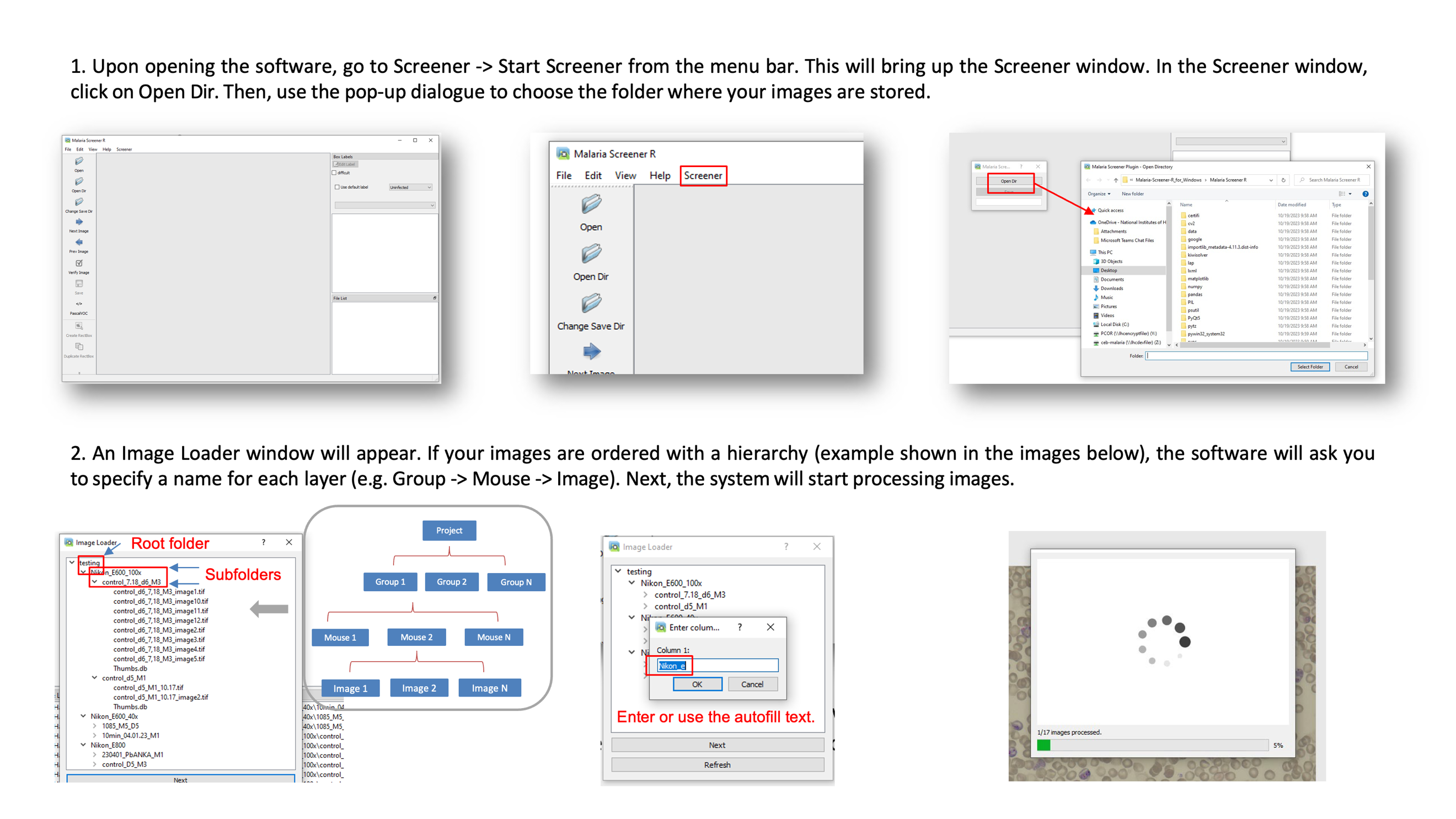

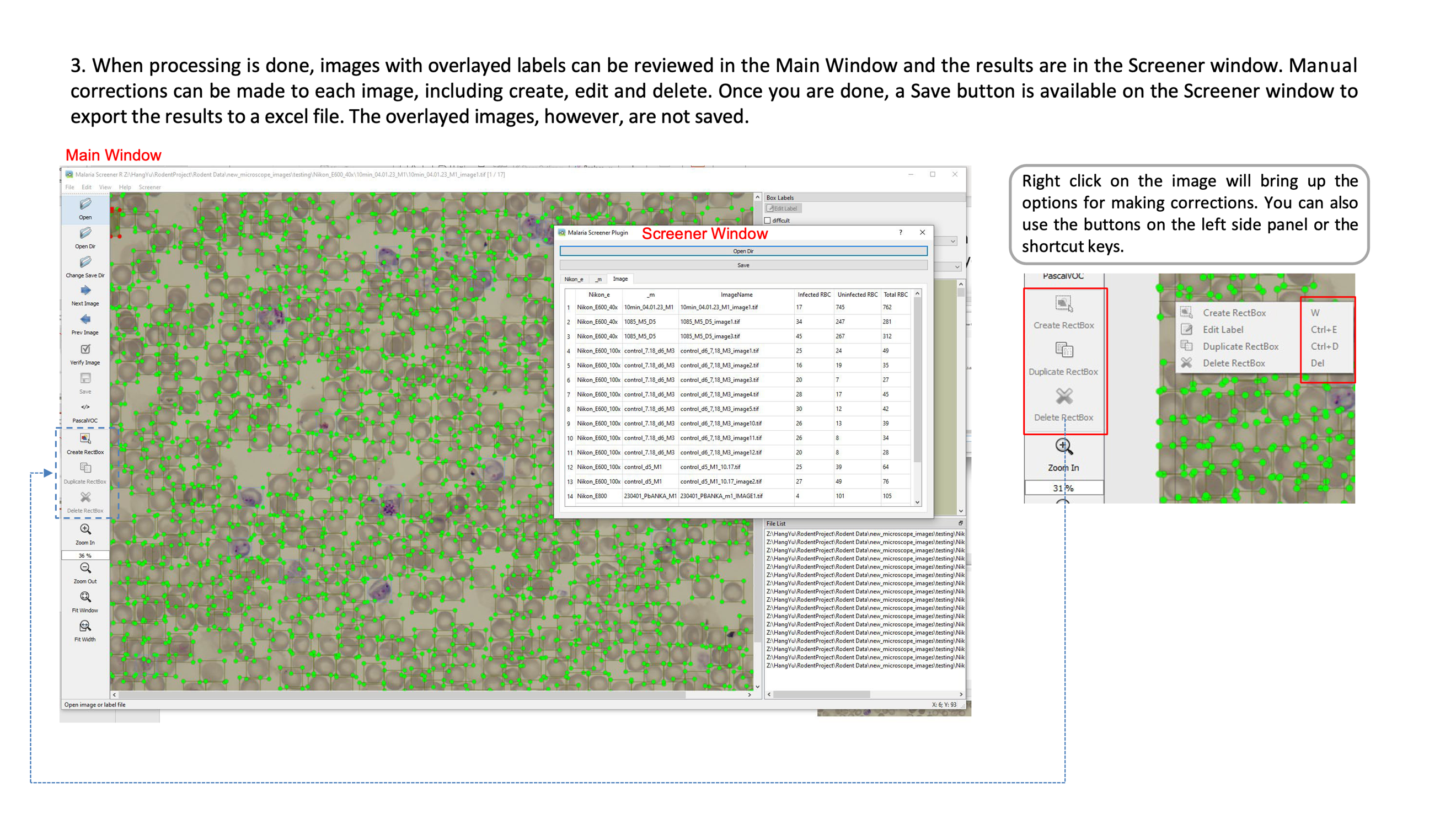

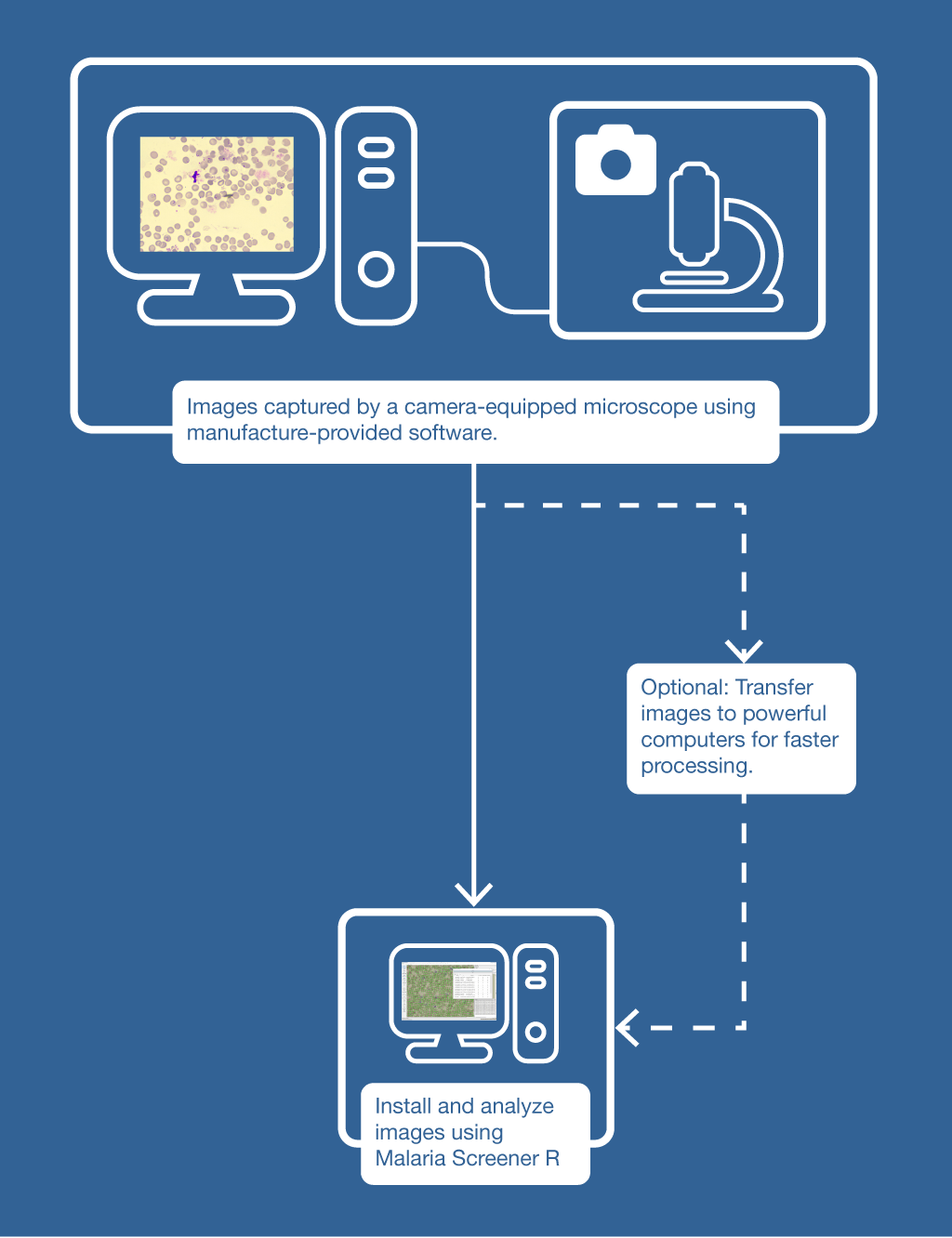

**Supplementary Figure 1: User manual demonstrating proper usage of software for automated detection of *Plasmodium* infected rodent RBCs.** A) A flow chart illustrates proper image capture and transfer computer with to Malaria Screener R. B) Steps 1-3 outline software usage for automated detection of RBCs from images captured under microscope.

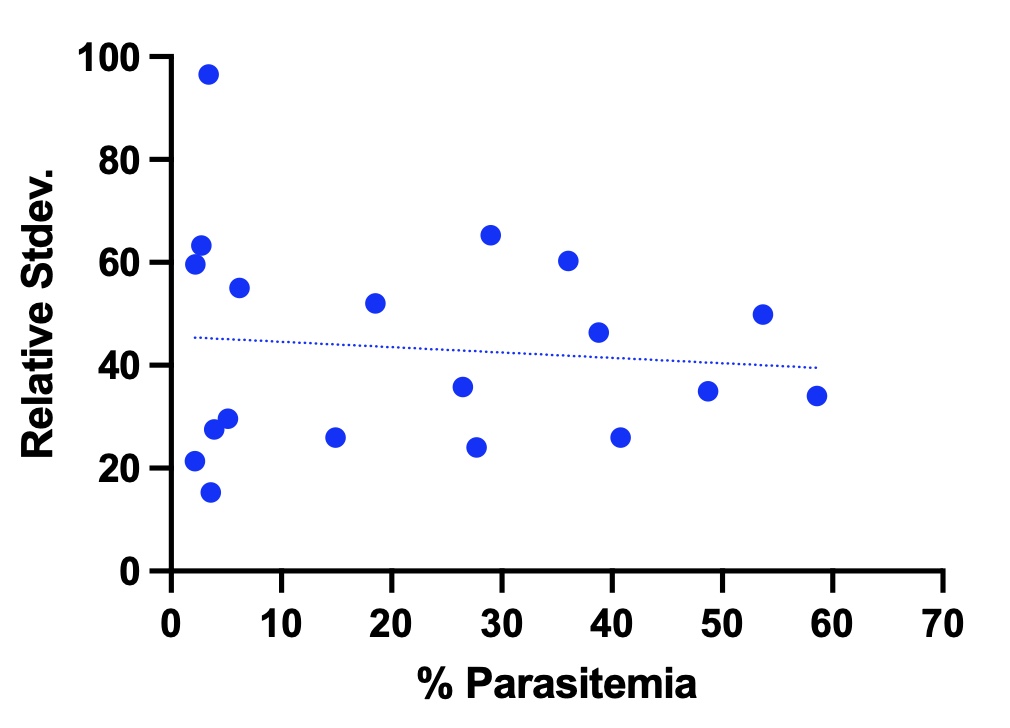

**R^2^ = 0.0099**

**Supplementary Figure 2: Relative Standard Deviation vs. Percent Parasitemia.**

Blue circles represent the standard deviation (y-axis) between parasitemia estimates from 4 different parasitologist across a range of parasitemia values (x-axis). Blue dotted line is a simple linear regression of all data points. R^2^ value is shown in the upper right.

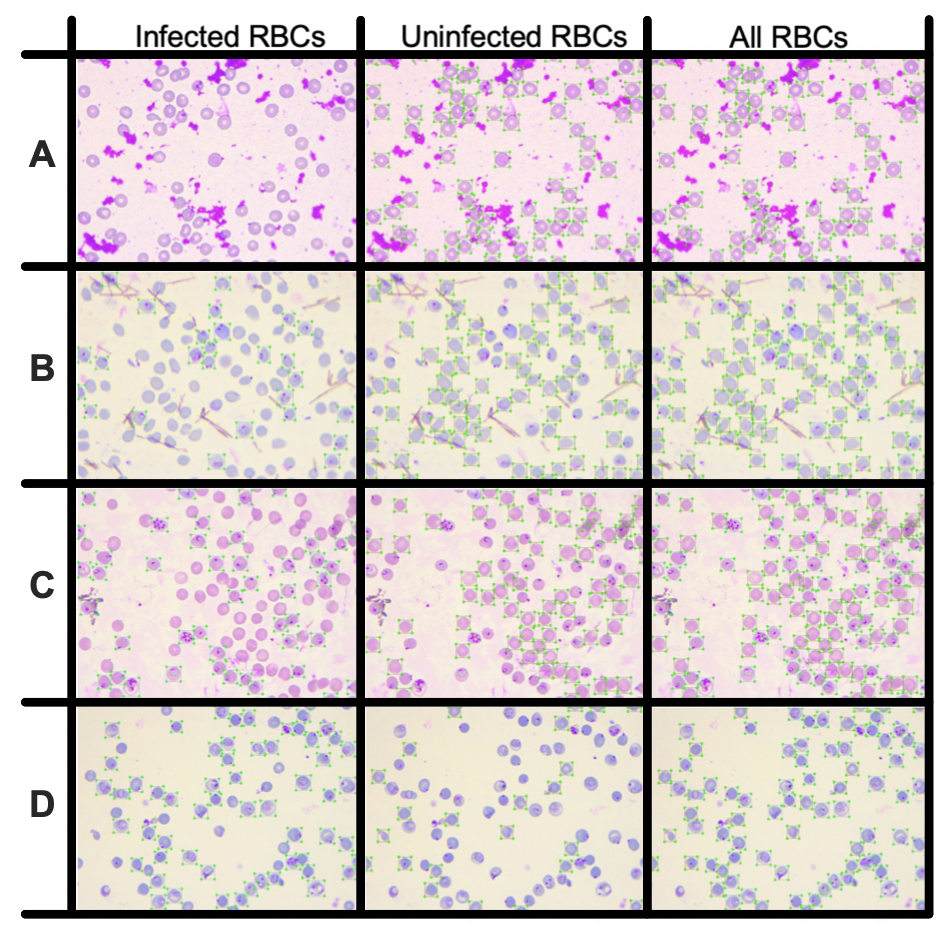

**Supplementary Figure 3: Example images of model performance on *P. yoelii* infected rodent RBCs.** Four example images of the model’s (Model_Rodent_RBCs) automated detections are shown at 0 % (A), 25% (B), 39% (C), and 68% (D) parasitemia respectively. Left column highlights RBCs labeled by the model as infected. Center column highlights RBCs labeled by the model as uninfected. Right column highlights all detected RBCs by the model. Green circumscribing boxes represent positive labels for each category.

**A**

**Blood Smears**

**Images**

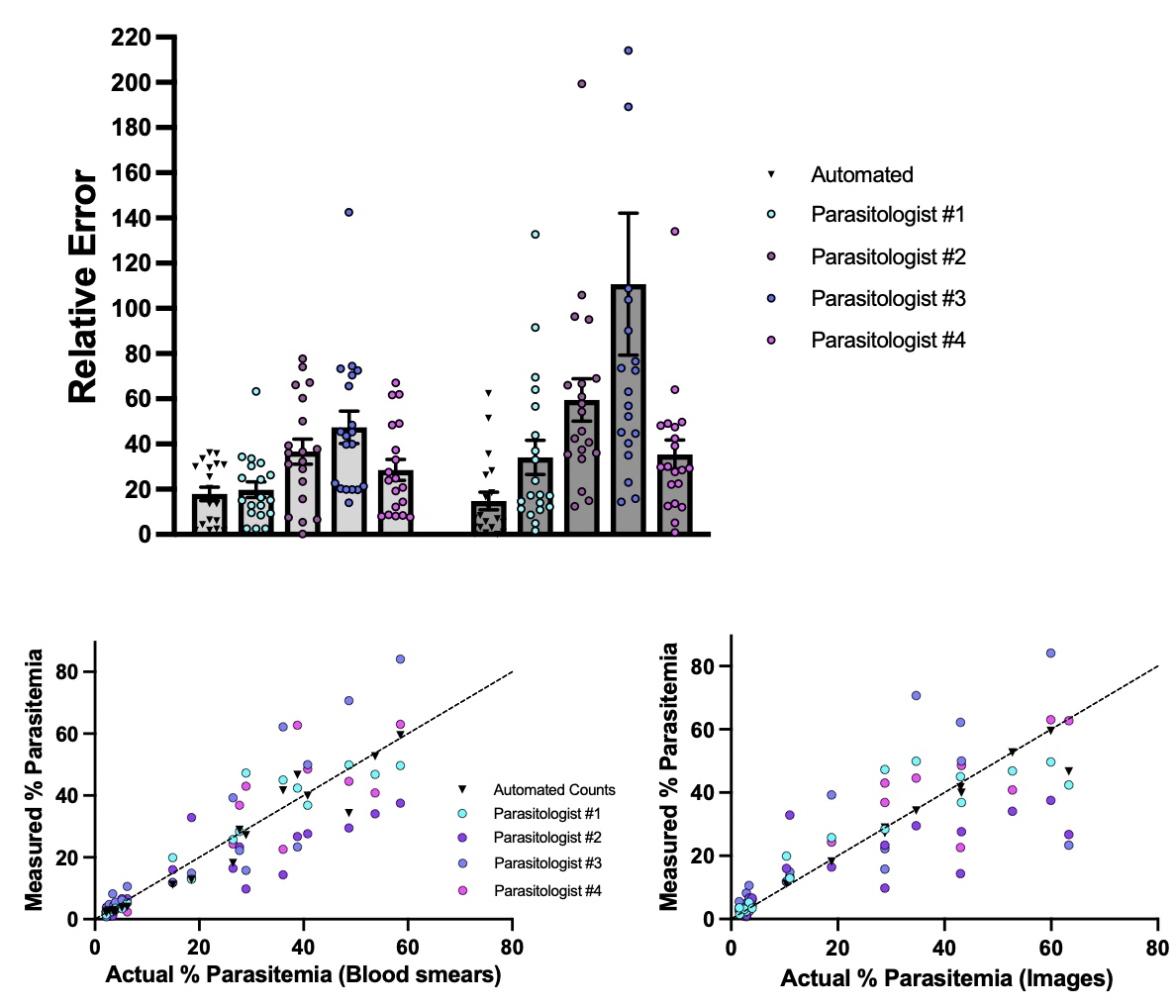

**B**

**C**

**Supplementary Figure 4: Relative Error of Automated Method vs. Individual Parasitologists.** A) Relative error values are shown for a single user determining % parasitemia by capturing images of blood smears and analyzing them using the automated model, Model_Rodent_RBCs (Black triangles), or four parasitologists manually counting infected/uninfected RBCs on the same blood smears under a microscope (Par. #1 = Cyan, Par. #2 = Purple, Par. #3 = Blue, and Par. #4 = Magenta). Relative error is calculated for each blood smear as the absolute value of ((% parasitemia user – % *parasitemia reference standard*)/(% *parasitemia reference standard*)*100). *% parasitemia reference standard* was calculated using two different methods, by taking the mean % parasitemia of manual counts of the blood smears by 4 expert parasitologists (left) or by taking the % parasitemia counted by a single parasitologist counting every cell on images captured from the corresponding blood smears (right). B-C) % Parasitemia (y-axis) measured by capturing images of blood smears and analyzing them using the automated model (Model_Rodent_RBCs) (Black triangles) or four parasitologists manually counting infected/uninfected RBCs on the same blood smears under a microscope (Par. #1 = Cyan, Par. #2 = Purple, Par. #3 = Blue, and Par. #4 = Magenta). X-axis corresponds to the % parasitemia calculated using (B) the average of manual estimates of blood smears by all 4 parasitologists or from (C) a single parasitologist individually counting each cell in the images taken of the same blood smears.

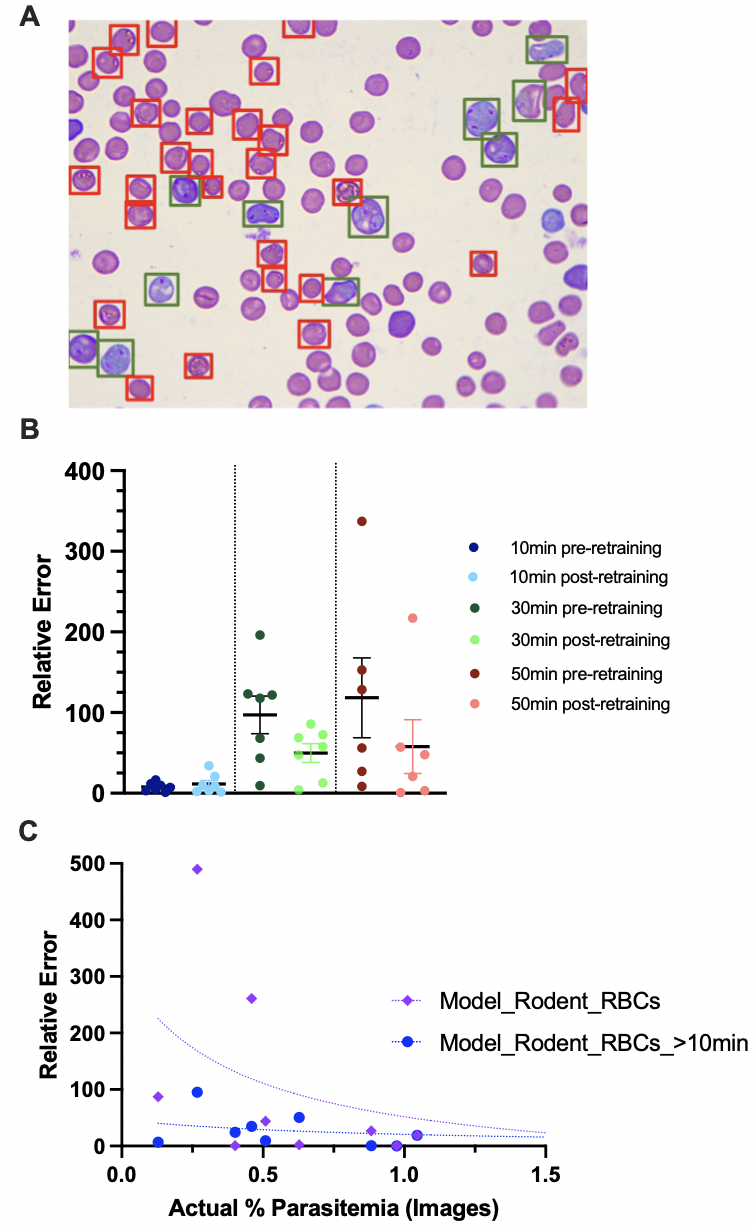

**Supplementary Figure 5: Effect of Giemsa stain time on model performance.** A) Example image is shown of a blood smear from a *P. berghei* infected mouse stained with Giemsa for 70 minutes. All RBCs annotated as infected by the original model (Model_Rodent_RBCs) are boxed. RBCs labeled as uninfected by the model are left unboxed. Green boxes represent infected RBCs that are true positives, and red boxes represent infected RBCs that are false positives. B) Relative error values are shown comparing detections by Model_Rodent_RBCs vs. manual counts using images from blood smears stained for 10min (dark blue, left), 30min (dark green, center), and 50min (dark red, right). Relative error is calculated for each mouse at one time point as the absolute value of ((% parasitemia automated count – % parasitemia manual count)/(% parasitemia manual count)*100). The model was then retrained with additional images from blood smears stained for 20min or 70min. This updated model (Model_Rodent_RBCs_>10min) was used to annotate the same images. Relative error using this updated model is shown to the right of original data points in light blue, light green, and light red respectively. C) Relative error of automated parasitemia measurements of *P. yoelii*-infected mice is shown from 0-6% parasitemia. Purple and blue dots represent relative error between automated and manual measurements for 1 mouse at 1 time point for the original and retrained model respectively. Relative error is calculated as the absolute value of ((% parasitemia automated count – % parasitemia manual count)/(% parasitemia manual count)*100). Purple and blue dotted lines represent nonlinear fit of relative error vs. % parasitemia for the original and retrained models respectively.
